# Codon usage bias in radioresistant bacteria

**DOI:** 10.1101/2020.02.25.964916

**Authors:** Maddalena Dilucca, Athanasia Pavlopoulou, Alexandros G. Georgakilas, Andrea Giansanti

## Abstract

The relationship between patterns of codon usage bias (CUB), the preferential usage of synonimous nucleotide triplets encoding the same amino acid, and radioresistance was investigated int he genomes of 16 taxonomically distinct radioresistant prokaryotic organisms and in a control set of 11 non-radioresistant bacteria. The radioresistant species were found to be strongly biased towards G and C in the third synonimous codon position. ENC and neutrality plots also sugest that CUB in radioresistant bacteria is mainly affected by mutational bias. Furthermore, the availability of tRNA gene copy number was analyzed and it was found that nine radioresistant species have the sam number of tRNA gene copies for each codon. This suggests that tRNA gene copies and codon bias co-evolved in a specific way in radioresistant species.

## 1 Introduction

Certain prokaryotic organisms, such as *Deinococcus radiodurans*, exhibit resistance to extremely high doses of ionizing radiation (Krisko and Radman, 2013; Makarova et al, 2001). Radioresistance and radiosensitivity (non-radioresistance) of prokaryotes could be attributed in part to specific evolutionary pressures upon these organisms, including the stress exerted by their host environment (Chanal et al, 2006). Of importance, codon usage bias (CUB), the unequal occurrence of synonymous codons in the coding sequences, was shown to differ among and within organisms (Plotkin and Kudla, 2011; Sharp et al, 2010). The preference of using a particular synonymous codon over another is affected to a great extent by environmental factors (Dittmar et al, 2006; Goodarzi et al, 2008), translation optimization (Rocha, 2004), tRNA abundance (Kahali et al, 2007), nucleotide composition (Choudhury et al, 2017; Novembre, 2002), protein structure (Oresic et al, 2003), protein length (Ingvarsson, 2007) etc. CUB reflects the dialectics between mutational bias (which generates codon diversity) and natural (Darwinian) selection against non optimal codons, which reduces codon diversity. In particular, mutations in the third codon position can affect an organism’s tRNA gene pool of the organisms and, conversely, the relative abundance of the gene copy numbers for isoacceptor tRNAs (tRNAs with different anticodons which carry the same amino acid) correlates with codon usage (Behura and Severson, 2011; Novoa et al, 2012). The theme of CUB and radioresistance seems to be still quite overlooked. Herein, we have studied the relationships between CUB and radioresistance in prokaryotes. We have selected a set of 16 radioresistant bacteria (comprising four species that were previously investigated by Pavlopoulou and colleagues (Pavlopoulou et al, 2016)) from the RadioP1 database and a set of known non-radioresistant prokaryotic organisms from diverse taxonomic divisions. We have compared CUB patterns in the two groups looking at GC contents, effective numbers of codons (ENC), relative codon usages, tRNA gene pools, ENC and neutrality plots. We found that radioresistant genomes are GC-rich, that their CUB is specifically tuned with tRNA availability, and ruled by mutational bias.

## 2 Materials and Methods

### 2.1 Genomes

The selection of the prokaryotic radioresistant organisms was based on a previous study by (Pavlopoulou et al, 2016) and on the RadioP1 database (http://doi.org/10.5772/60471). Only the most adequately annotated genomes of these organisms were downloaded from NCBI’s GenBank (Benson et al, 2017). tRNA data were downloaded from GtRNA (http://gtrnadb2009.ucsc.edu/).

In our study, we have also included a control group of eleven prokaryotes from different taxonomic divisions and environmental conditions, in order to compare their CUB patterns with the ones of the above organisms (Table 2).

**Table 1.** Radioresistant genomes. The species name, species abbreviation, class and NCBI RefSeq accession code. The radioresistant considered in (Pavlopoulou et al, 2016) are in bold. Classes: *Actinobacteria* (1), *Alphaproteobacteria* (2), *Deinococci* (3), *Gloebacteria* (4),*Halobacteria* (5), *Thermococci* (6).

| Organism | Abbreviation | Class | Accession code |
| --- | --- | --- | --- |
| *Deinococcus deserti* VCD 115 | dede | 3 | CP001114.1 |
| *Deinococcus geothermalis* DSM 11300 | dege | 3 | CP000359.1 |
| *Deinococcus gobiensis* I-0 | dego | 3 | CP002191.1 |
| *Deinococcus maricopensis* DSM 21211 | dema | 3 | CP002454.1 |
| *Deinococcus proteolyticus* MRP | depr | 3 | CP002536.1 |
| *Deinococcus radiodurans* R1 | **dera** | 3 | NC001263.1 |
| *Geodermatophilus obscurus* DSM 43160 | geob | 1 | CP001867.1 |
| *Gloeobacter violaceus* PCC 7421 | glvi | 4 | BA000045.2 |
| *Halobacterium salinarum* R1 | **hasa** | 5 | NC010364 |
| *Haloferax volcanii* DS2 | havo | 5 | CP001956.1 |
| *Kineococcus radiotolerans* SRS30216 | **kira** | 1 | NC009664 |
| *Methylobacterium radiotolerans* JCM 2831 | mera | 2 | CP001001.1 |
| *Pyrococcus furiosus* DSM 3638 | pyfr | 6 | AE009950.1 |
| *Pyrococcus furiosus* COM1 | pyfu | 6 | CP003685.1 |
| *Thermococcus gammatolerans* EJ3 | **thga** | 6 | CP001398 |
| *Truepera radiovitrix* DSM 17093 | trra | 3 | CP002049.1 |

**Table 2.** Non-radioresistant bacteriall genomes. The species name, species abbreviation, class and NCBI RefSeq accession code. Classes: *Actinobacteria* (1), *Alphaproteobacteria* (2), *Bacilli* (3),*Bacteroidetes* (4), *Betaproteobacteria* (5), *Chlamydiae* (6), *Deltaproteobacteria* (7), *Epsilonproteobacteria* (8), *Gammaproteobacteria* (9),*Mollicutes* (10), *Spirochaetes* (11).

| Organism | Abbreviation | Class | Accession code |
| --- | --- | --- | --- |
| *Agrobacterium tumefaciens* (fabrum) | agtu | 2 | NC003062 |
| *Bacteroides thetaiotaomicron* VPI-5482 | bath | 4 | NC004663 |
| *Bdellovibrio bacteriovorus* HD100 | bdba | 7 | BX842601 |
| *Burkholderia pseudomallei* K96243 | bups | 5 | NC006350 |
| *Chlamydia trachomatis* D/UW-3/CX | chtr | 6 | NC000117 |
| *Helicobacter pylori* 26695 | hepy | 8 | NC000915 |
| *Mycoplasma genitalium* G37 | myge | 10 | NC000908 |
| *Mycobacterium tuberculosis* H37R | mytu | 1 | NC009525 |
| *Staphylococcus aureus* N315 | stau | 3 | NC002745 |
| *Treponema pallidum* Nichols | trpa | 11 | NC000919 |
| *Vibrio cholerae* N16961 | vich | 9 | NC002505 |

### 2.2 Codon bias indices

#### ENC (Effective Number of Codons)

(Wright, 1990) *ENC* is an estimate of the frequency of different codons used in a coding sequence; it expresses the codon bias of a gene by making quantitative its tendency to use a restricted set of codons. In principle *ENC* ranges from 20 (when each aminoacid is coded by just one and the same codon) to 61 (when all synonymous codons are used on an equal footing). Given a sequence of interest, the computation of *ENC* starts from *F*_*α*_, a quantity defined for each family *α* of synonymous codons (one for each amino acid):

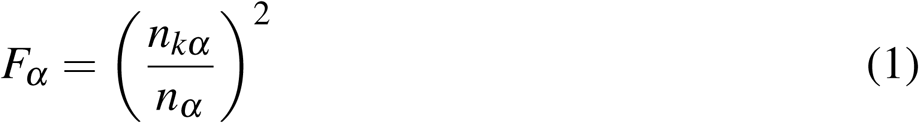

where *m*_*α*_ is the number of different codons in *α* (each one appearing 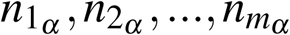 times in the sequence) and 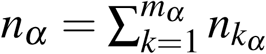.

*ENC* then weights these quantities on a sequence:

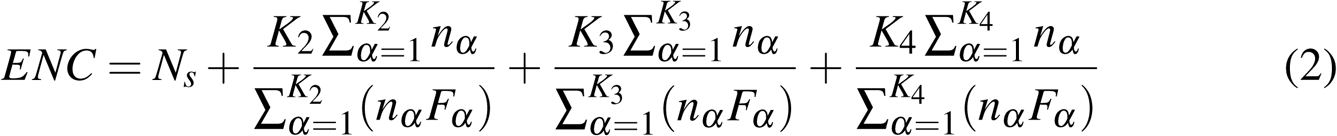

where *N*_*S*_ is the number of families with one codon only and *K*_*m*_ is the number of families with degeneracy *m* (the set of 6 synonymous codons for *Leu* can be split into one family with degeneracy 2, similar to that of *Phe*, and one family with degeneracy 4, similar to that, e.g., of *Pro*).

In this paper we have evaluated *ENC* by using the implementation in *DAMBE* 5.0 (Xia, 2013).

#### GC content

In genetics, the *GC* content is the percentage of the nitrogenous bases on a DNA molecule that are either guanine or cytosine. The overall *GC* content, especially *GC*_3_ (i.e. the GC content at the third codon position), frequently reflects the strength of mutational bias. The *GC* content at the first *GC*_1_ second *GC*_2_, and third *GC*_3_codon position was computed using a PERL script from the inspiring study on microsporidia by (Xiang et al, 2015) (https://github.com/hxiang1019/calcGCcontent.git). The three stop codons (UAA, UAG, and UGA) and the three codons encoding the amino acid residue isoleucine (AUU, AUC, and AUA) were excluded from the calculation of *GC*_3_, as well as the single codons corresponding to methionine (AUG) and tryptophan (UGG) were excluded from the calculation of *GC*_1_, *GC*_2_, and *GC*_3_.

To test whether the 67 radioresistance signature genes proposed in (Pavlopoulou et al, 2016) display peculiar ENC and GC content values, the *Z*_*score*_ values for each gene was calculated as:

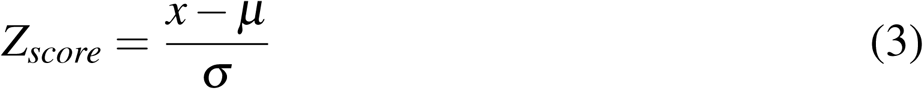

where x is, respectively, the codon bias index of a gene (*ENC*) and then the *GC* content, *μ* is the average value of the indexes over all the genes of the species and *σ* the standard deviations. Genes with *Z*_*score*_ values above 1 were considered to have higher codon bias than the mean value. These genes are shown in red in the Excel file in Supplementary Materials.

#### RSCU (Relative synonymous codon usage)

*RSCU* vectors for all the genomes were computed by using the implementation in DAMBE 5.0 (Xia, 2013), following the formula:

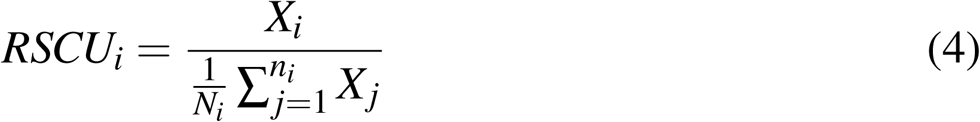

In the *RSCU*_*i*_ *X*_*i*_ is the number of occurrences, in a given genome, of codon i, and the sum in the denominator runs over its *n*_*i*_ synonymous codons; *RSCU*_*s*_ then measure codon usage bias within a family of synonymous codons. If the *RSCU* value for a codon *i* is equal to 1, this codon was chosen equally and randomly. Codons with *RSCU* values greater than 1 have positive codon usage bias, while those with value less than 1 have relatively negative codon usage bias (Sharp and Li, 1986). *RSCU* heat maps were drawn with the CIMminer software (Weinstein et al, 1997), which uses Euclidean distances and the average linkage algorithm.

### 2.3 ENC plot

The ENC-plot is a well known tool to investigate the patterns of synonymous codon usage in which the *ENC* values are plotted against *GC*_3_ values when codon usage is dominated by the mutational bias, the formula of expected *ENC* values is given by:

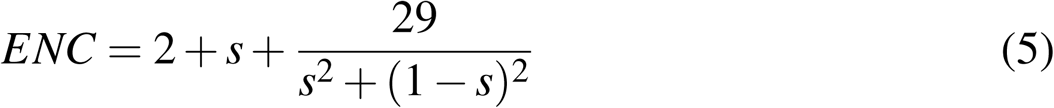

where *s* represents the value of *GC*_3_ (Wright, 1990). When the corresponding points fall near the expected neutral curve, mutations that enforce the typical mutational bias of the species are the main factor affecting the observed codon diversity. Whereas when the corresponding points fall considerably below the expected curve, the observed CUB is mainly affected by natural selection.

### 2.4 Neutrality plot

Neutrality plots were drawn by plotting *GC*_12_, the average value of *GC*_1_ and *GC*_2_ on the vertical axis and *GC*_3_ on the horizontal axis; as in the ENC plot each dot represents an independent gene. If the slope of the regression line is close to 1, there is statistically significant correlation between *GC*_12_ and *GC*_3_, pointing to a codon usage due to neutral mutational bias. Conversely, lack of correlation between *GC*_12_ and *GC*_3_ indicates that the whole genome is under selective pressure (Sueoka, 1999).

## 3 Results

### Genomes of radioresistant species are GC enriched

The *GC* distribution over the genes of radioresistant and non-radioresistant species is shown in Figure 1. Radioresistant species have an overall *GC* content significantly higher than 50%, whereas non-radioresistant bacteria display a rather bimodal distribution. The two distributions are significantly distinct (Mann-Whitney-Wilcoxon U test, p-value ≤0.5). This shows that G/C-ending codons are systematically preferred over A/U-ending ones in radioresistant organisms. Suggesting that radioresistant genomes have acquired a relatively higher thermal stability. Moreover, in radioresistant species *GC*_3_ has the highest mean value of 81.63 % (SD=1.63), followed by *GC*_1_ (mean= 68.60%, SD 0.9) and *GC*_2_ (mean=58.49%, SD=1). In non-radioresistant species, *GC*_1_ has the highest mean value of 58.49% (SD=1), followed by *GC*_3_ (mean= 56.20%, SD 2.2) and *GC*_2_ (mean=40.94%, SD=0.8) (see box plots of Figure 2). It has been previously shown that GC content and CUB are correlated (Li et al, 2015). Given that the genomic *GC* content is an important evolutionary signature (Šmarda et al, 2014), (Lassalle et al, 2015), the above observation suggests that radioresistance and CUB should be correlated. Indeed, as shown above, *GC*_3_ in radioresistant species is significantly higher than the average *GC* content, suggesting that in these species CUB plays a more relevant role than in non-radioresistant species.

**Fig. 1.**
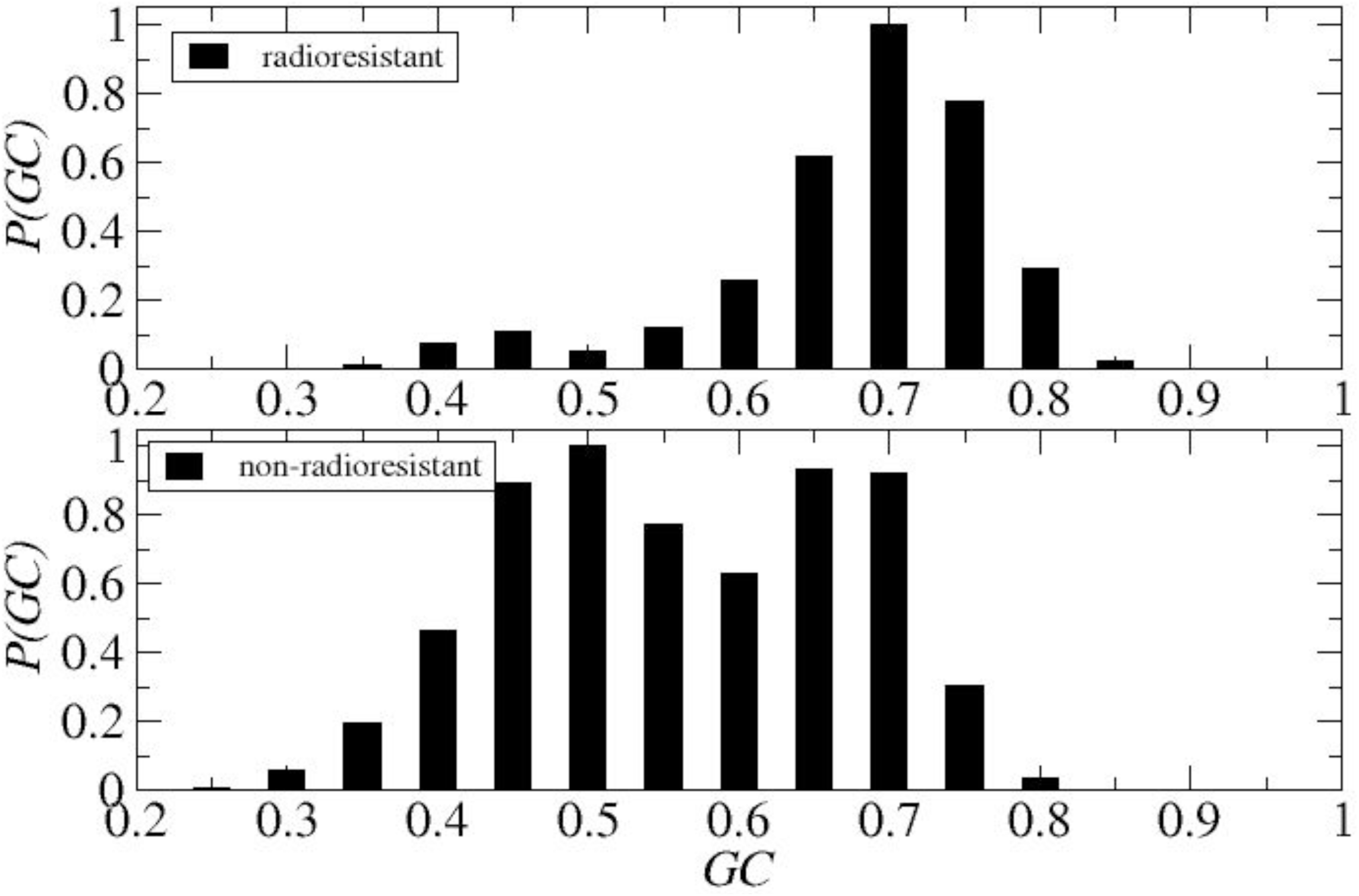
Distribution of GC contents. The GC distributions of radioresistant and radiosensitive species are significantly different: average values are 0.66 ± 0.09 (top panel) and 0.53 ± 0.11 (bottom panel) respectively (Mann-Whitney-Wilcoxon U test, p-value ≤ 0.5).

**Fig. 2.**
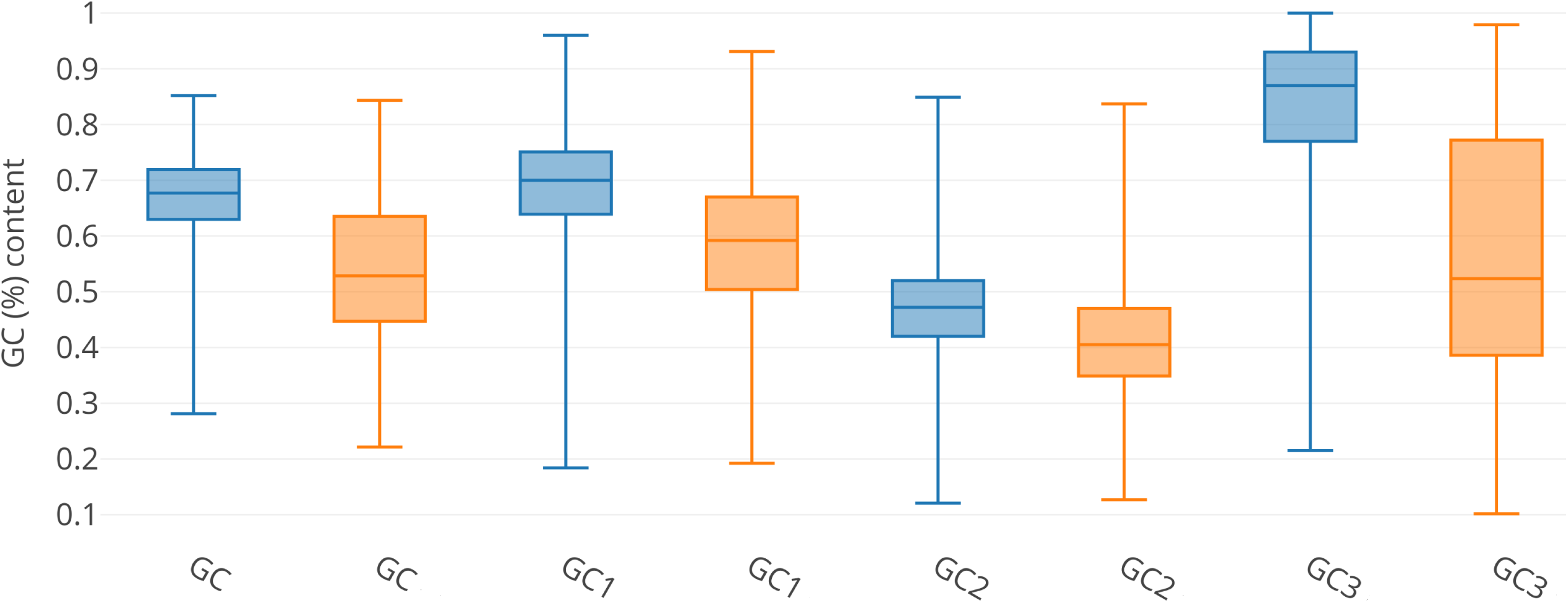
GC contents. Box plot of GC content variation in different coding position. Blue boxes: radioresistent species. Orange boxes: non-radioresistant species (the bottom and the top of the box are the lower and upper quartiles, respectively, and the end of the whiskers are, respectively, the lowest and the highest data still within 1.5 times the interquartile range of the lower and higher quartiles.)

### Codon usage bias

Indeed, figure 3 shows that radioresistant species have effective numbers of codons signifcantly different from those of non-radioresistant species. In particular, radioresistant species use a more restricted repertoire of codons with respect to that of non-radioresistant ones.

**Fig. 3.**
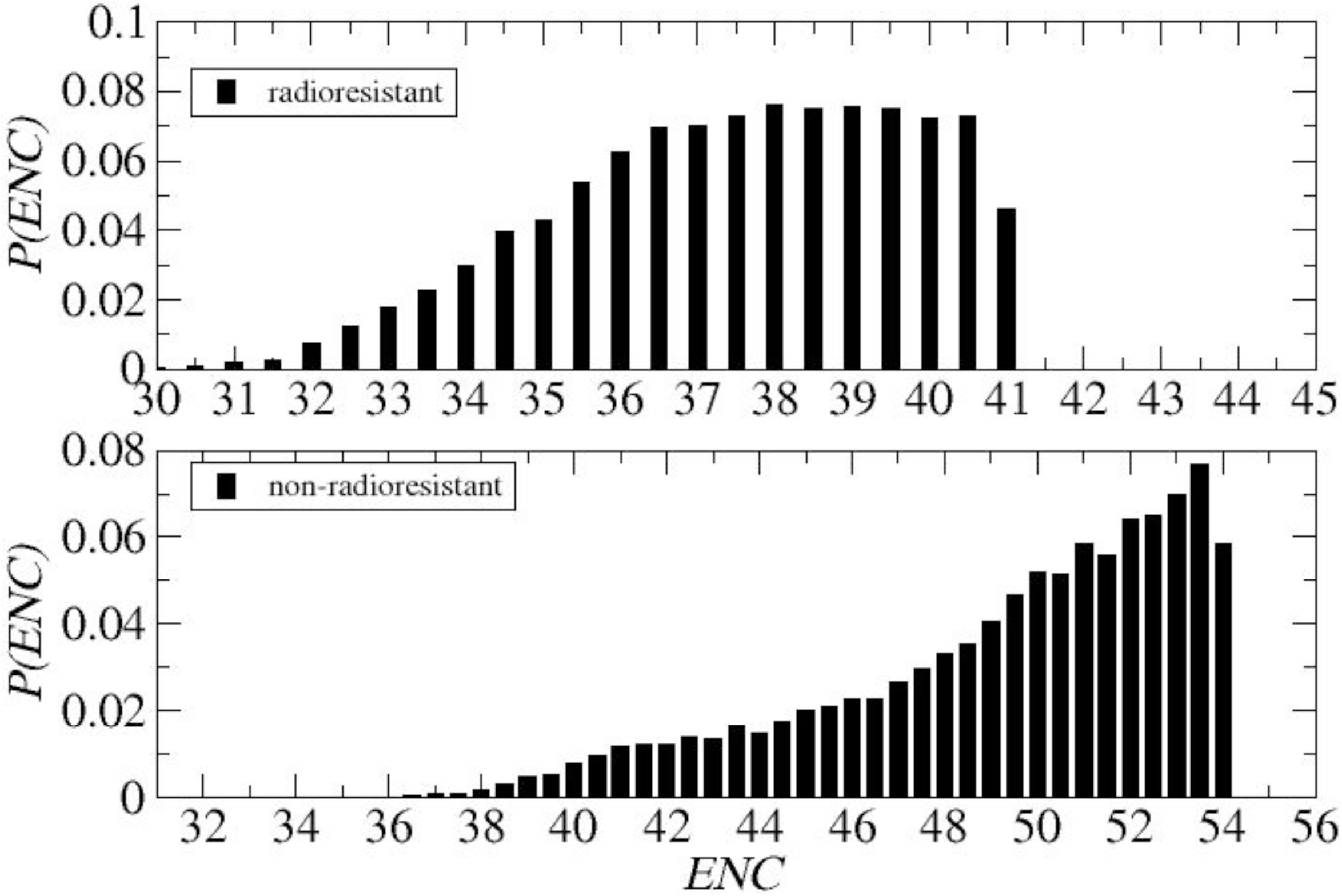
ENC distributions.

Codon usage patterns in the radioresistant genomes were investigated also by evaluating Relative Synonymous Codon Usages (*RSCU* values). As it is shown in Figure 4, radioresistant species evolved to code their amino acids using a restricted set of about optimal (G/C-ending) codons. Since the non-radioresistant species in this heat map where chosen as representative of several taxonomical classes this observation strongly suggests that adaptation to intense radiation induces a specific response in the codon bias of the radioresistant species.

**Fig. 4.**
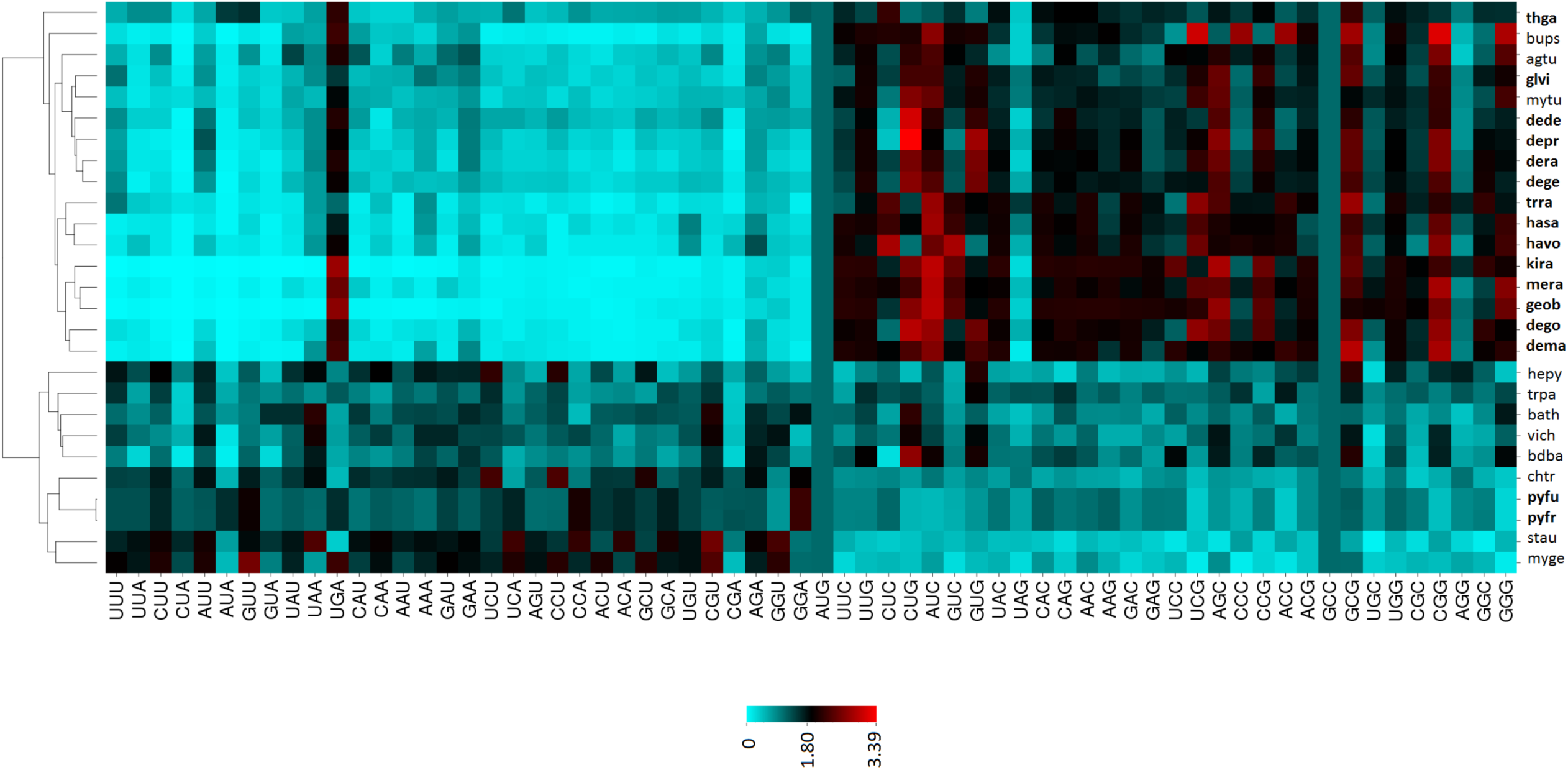
Heatmap of RSCU values. The heatmap was created with CIMminer. Greater RSCU values, indicating preferential codon usage, are represented by darker shades of red. The radioresistant bacteria strongly prefer G/C-ending codons and form a distinct cluster as compared to the control group of bacteria. Radioresistant bacteria are written in bold.

The heatmap of RSCUs confirms that the radioresistant species are sistematically enriched in GC-ending codons (see Figure 4, upper right) sustained by high tRNA availability (see Figure 5, upper right). This is at variance with what is observed in non-radioresistant species, that sistematically incorporate AU-ending codons, that is sustained by enrichment of the corresponding tRNAs. These observations clearly indicate that the emergence of radioresistance rests on the co-evolution of CUB and tRNA availability. A co-evolution that has generated codon and amino-acid distributions which are significantly different in radioresistant and non-radioresistant species. It is worth nothing that the combined use of the heat maps in Figures 4 and 5 could provide a CUB signature usefull to search, among bacterial genomes, for species that could emerge as radioresistant, under radioactive stress.

**Fig. 5.**
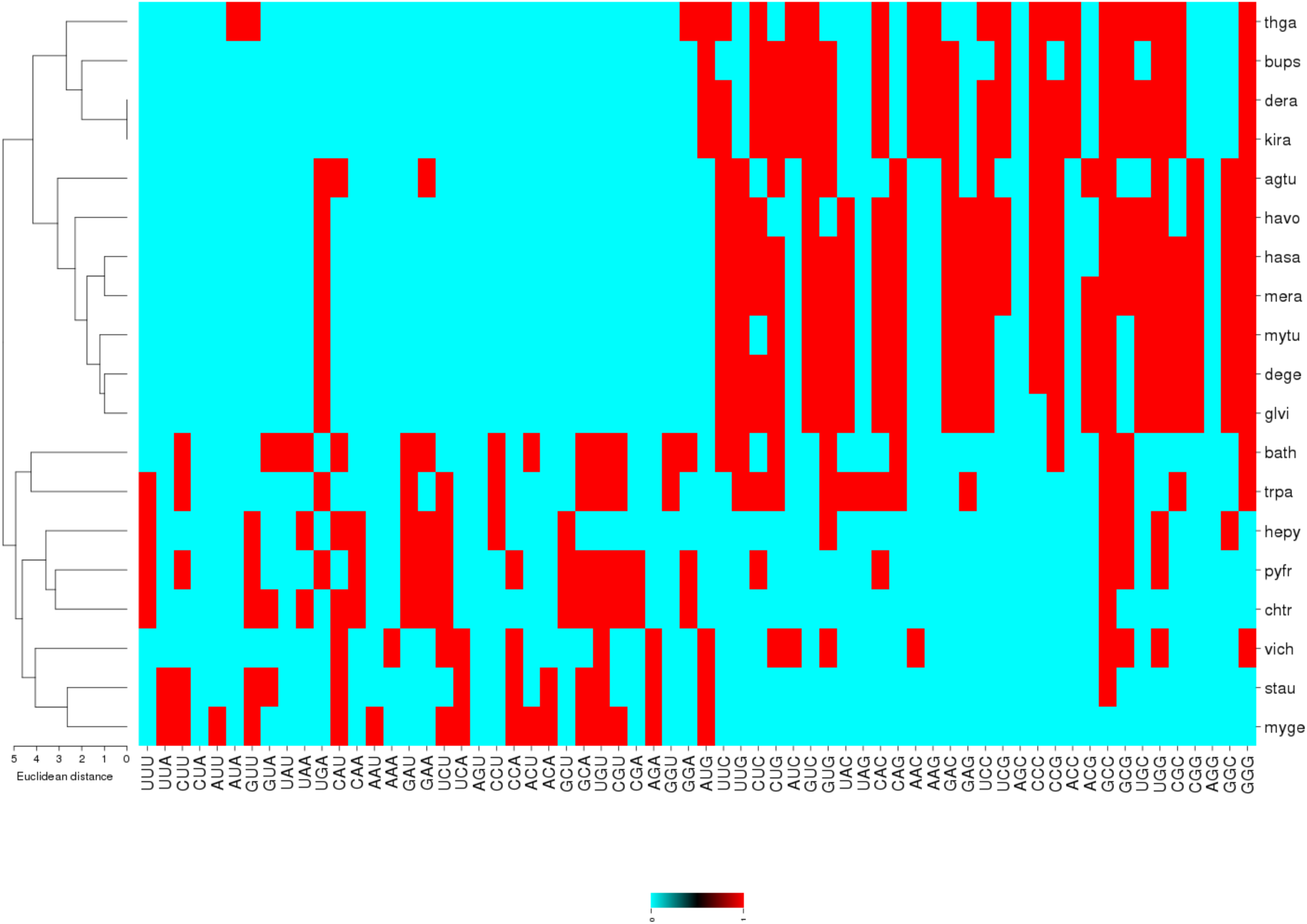
Heatmap of tRNA values.

### ENC - *GC*_3_ plots

As specified in the methods *ENC* vs *GC*_3_ plots and neutrality plots are first-choice tools to quantitatively measure how much the choice of codons in a gene is either under mutational bias or natural selection. Separate *ENC* plots, for radio resistant and non-radioresistant species are reported in Figure 6, where it is visually evident that the representative centroids of radioresistant species are closer to the solid curve which encods the pure case of mutational bias. Non-radioresistant species appear, at least in several cases, more prone to have their codon bias affected by natural selection. To make more quantitative this observation in Figure 7 the average distances from the reference curve (See Equation 2.3) in the ENC plot are reported, separating the genes of radioresistant from those of non-radioresistant species. Genes were also separated into COGs, clusters of othologous genes (Tatusov et al, 2001), to investigate the possible dependence of evolutionary pressures on functional specialisation. Interestingly, it is evident that on the average the genes of radioresistant species are significantly closer to the curve, pointing to a dominance of mutational bias. Moreover, it is evident that the different character of the evolutionary pressure exerted on the genes from radioresistant and non-radioresistant species is systematic, independent on the COG class. A consistent result was obtained also considering neutrality plots. In Figure 8 centroids of the different species considered here are represented on the *GC*_3_vs*GC*_12_ plane (neutrality plot) where it is quite evident that the genes of radio resistant species display a correlation-regression line with a slope close to 1 point to the fact that genes of radioresistant species let emerge their codon bias under a stronger mutational bias. This is a way to express the fact that the genes of radioresistant species are constrained in their codon usage more by the necessity of keeping a remarkably high *GC* content than as an effect of Darwinian selection. Let us stress that, from Figures 7, there is a remarkable separation in the *ENC* and neutrality plots, of radioresistant and non-radioresistant species. The codon bias of radioresistant species significantly emergense as a product of mutational bias, which is quite universal. It does depends of the functional specialization, as represented here by the COG’s clusters.

**Fig. 6.**
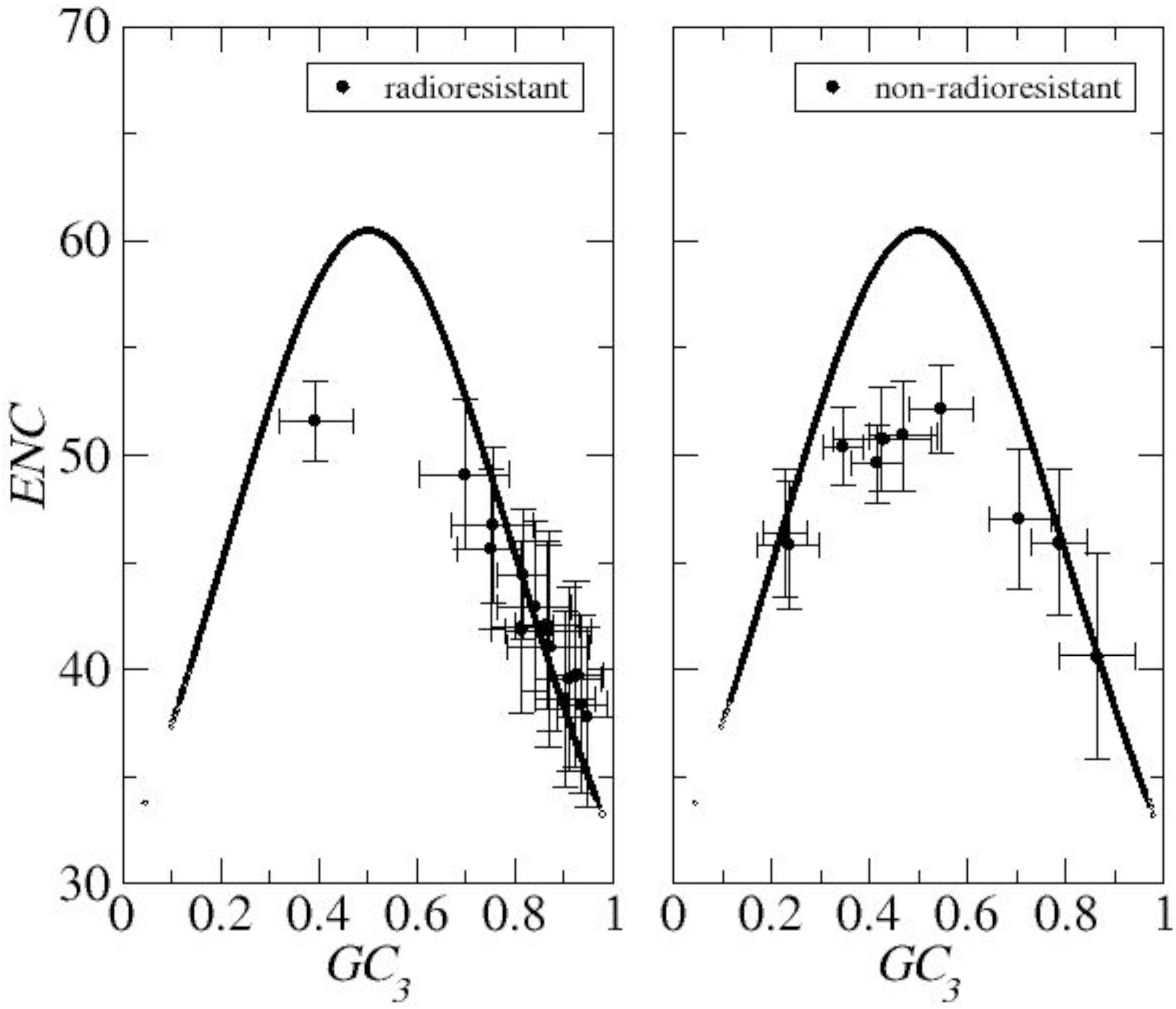
Centroids of ENC and *GC*_3_ for each bacteria.

**Fig. 7.**
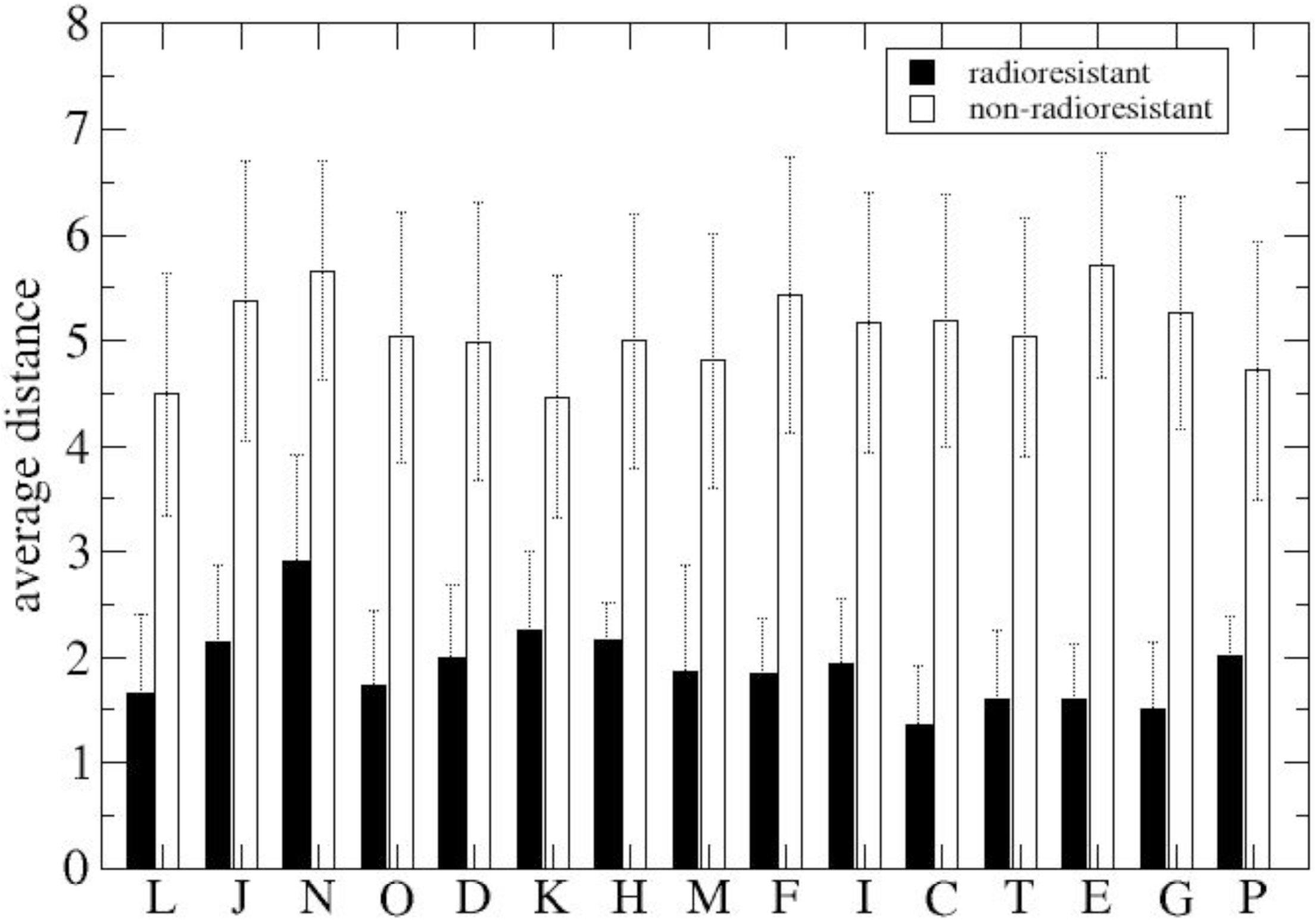
Distribution of distances of clusters of orthologous genes from solid curve *ENC*-*GC*_3_ plots. Note that radioresistant genes lie close to the curve.

**Fig. 8.**
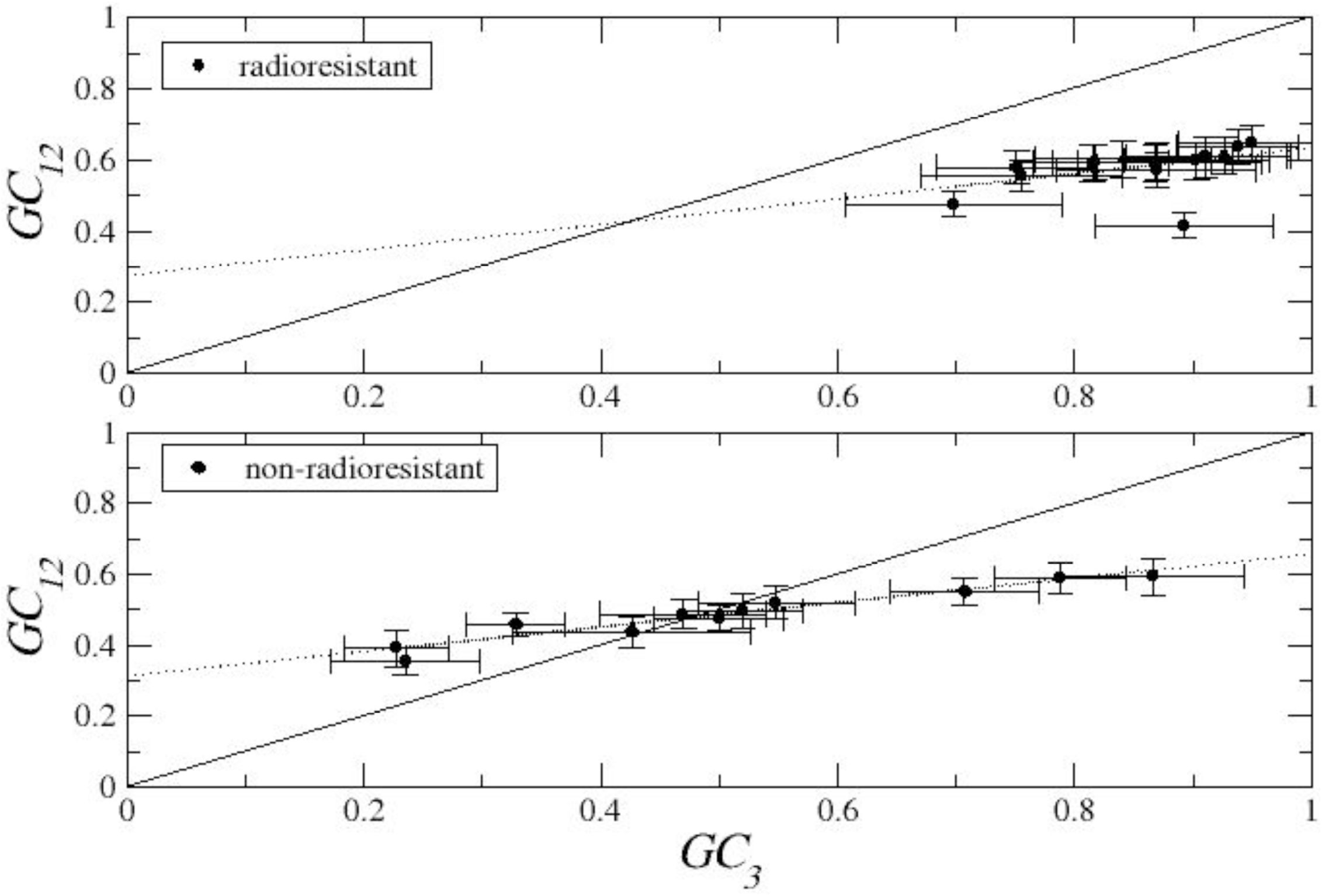
Neutrality plot for each bacteria.

### Network of radioresistance genes and CUB

As mentioned in the Introduction, Pavlopoulou et al. (see Table 2 in (Pavlopoulou et al, 2016)), recognized a core network of 67 homologous radioresistance genes (see also Supplementary Materials). Among these genes a restricted subset, namely: RecA, RuvB, DdrD, *TGAM_*200500, Lig, Mre11, RadB and UvrC, is strictly related to the emergence of radioresistance (their over-expression is a signature). These genes are mostly involved in the DNA damage repair machinery, which is the key process in the adaptation to ionizing and UV radiation. In the present context let us observe that all the above mentioned genes display a ENC *Z*_*score*_ bigger than 1. By abduction, we could suggest that most of the genes that are related to radioresistance display a strong CUB signature. Overall, these observations provide a strong indication for further genetic, evolutionary and engineering studies in the search for a universal network of genes related to the emergence of radioresistance. Interestingly, representing in ENC and neutrality plots, the genes in our dataset, that are homologous to the 67 genes selected in (Pavlopoulou et al, 2016), one observes that, consistently, they are under a significantly mutational bias (see Supplementary Materials).

## 4 Conclusions

Codon usage bias is a genomic fingerprint of a species, which basically reflects the adaptation of its translation mechanisms to environmental challenges. Radioresistant bacterial and archaeal species can resist to high doses of ionizing radiation. The relationship between codon bias and radioresistance is investigated an this study is one of the few which study. We have compared nucleotide composition (GC content) and patterns of codon usage in radioresistant species with respect to a control group of non-radioresistant bacteria. Moreover, we have investigated the codon bias of hubs of the protein-protein interaction networks associated to genes that are essential for radioresistance. The nucleotide composition analysis revealed that the overall GC content in radioresistant genomes is higher than 0.50, leading to the suggestion that G/C-ending codons might be preferred over A/U-ending codons (as then confirmed by RSCU analysis). It is generally known that highly expressed genes have a tendency to incorporate more Cs and Gs at synonymous positions than less expressed genes and that preference of C-ending codons is related to translational efficiency (Sharp and Li, 1987) and translational fidelity (Gouy and Gautier, 1982). Then, highly efficient mechanisms of gene expression seem to be necessary for a species to be (or to become) radioresistant. In fact, the results of correlation between GC content in the first two and the third position (*GC*_3_ and *GC*_12_) suggest that mutational bias may play an important role, particularly in radioresistant signature genes. This is a result that confirms what has been alredy observed in *D.radiodurans* by (Q., 2006). On the basis of the tRNA gene copy numbers, it can be inferred that organisms under ionizing radiation, may evolve specific tRNA gene pools (Novoa et al, 2012), thereby inducing codon usage bias. In particular, we observed that radioresistant bacteria, at variance with non-resistant species, display a specific repertoire of tRNA genes (see Figure 5). Synonymous codon usage at corresponding mRNA regions has been shown to affect the local structure and function of the encoded proteins (Saunders and Deane, 2010; Spencer and Barral, 2012). Although the amino acid composition of two polypeptide chains encoded by two mRNAs with synonymous codons is apparently the same, their corresponding proteins may adopt alternative conformations (Komar, 2007). For instance, the presence of a synonymous single nucleotide polymorphism (SNP) in the MDR1 (multidrug resistance 1) gene has been found to alter the substrate specificity of its product, P-glycoprotein (Kimchi-Sarfaty et al, 2007). Therefore, a protein with identical or very similar amino acid composition in two different organisms (i.e. radioresistant and non-radioresistant) can have different functions. This is the case of RecA in *E.Coli* e *D.Radiodurans*. The fact that most of the genes/proteins mentioned above (see Table 2 in (Pavlopoulou et al, 2016)) as participating in species-specific radioresistance networks leads to the suggestion that these molecules play a prominent role in radiotolerance in the respective species by being physically or functionally linked to other radioresistance-related proteins. We have clearly observed that the high connectivity of the radioresistance signature genes is definetly associated with highly biased codon usage.

## Author Contributions

MD AP AG AG designed the experiments; MD AG performed the experiments. MD AP collected data and wrote routines; ALL wrote and revise the paper. Conflicts of Interest: The authors declare no conflict of interest.

