## supplementary material for "Codon usage bias in radioresistant bacteria"

manuscript No.
(will be inserted by the editor)

### Codon usage bias in radioresistant bacteria
### SUPPORTING INFORMATION

**Maddalena Dilucca · Athanasia Pavlopoulou · Alexandros G. Georgakilas · Andrea Giansanti**

Received: date / Accepted: date

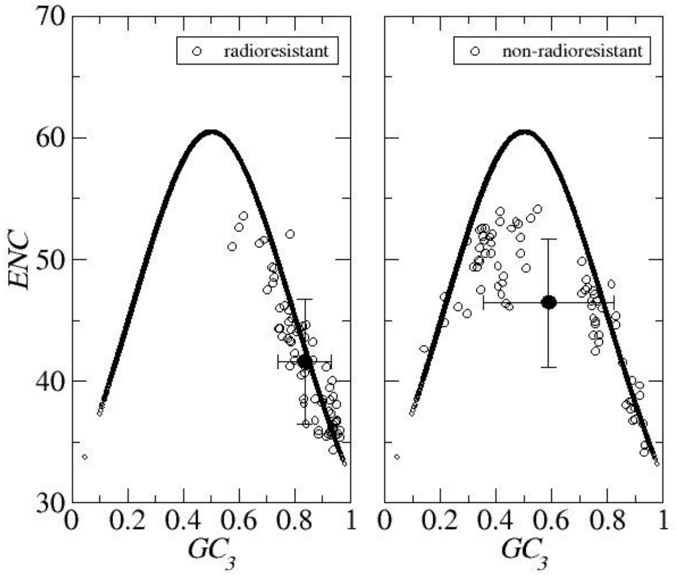


**Fig. 1 ENC-*GC*$_3$ plots of genes that are homologous to the 67 genes in Pavlopoulou et al.** Note that also in this subset the genes in radioresistant species are definitely closer to the solid curve.

Address(es) of author(s) should be given

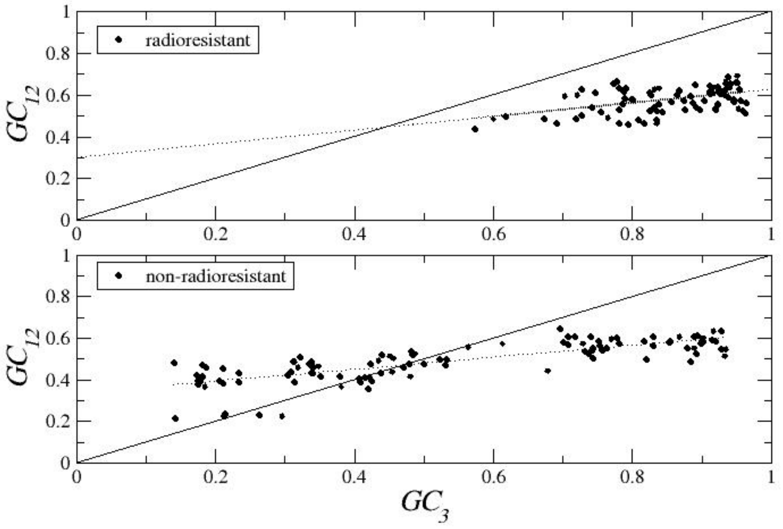


**Fig. 2 Neutrality plot of genes that are homologous to the 67 genes in Pavlopoulou et al.** Note that genes in radioresistant species lie far from bisector.
